# Two strategies underlying the trade-off of hepatitis C virus proliferation: stay-at-home or leaving-home?

**DOI:** 10.1101/821710

**Authors:** Shoya Iwanami, Kosaku Kitagawa, Yusuke Asai, Hirofumi Ohashi, Kazane Nishioka, Hisashi Inaba, Shinji Nakaoka, Takaji Wakita, Odo Diekmann, Shingo Iwami, Koichi Watashi

## Abstract

Viruses proliferate through both genome replication inside infected cells and transmission to new target cells or to new hosts. Each viral genome molecule in infected cells is used either for amplifying the intracellular genome as a template (“stay-at-home strategy”) or for packaging into progeny virions to be released extracellularly (“leaving-home strategy”). The balance between these strategies is important for both initial growth and transmission of viruses. In this study, we used hepatitis C virus (HCV) as a model system to study the functions of viral genomic RNA in both RNA replication in cells and in progeny virus assembly and release. Using viral infection assays combined with mathematical modelling, we characterized the dynamics of two different HCV strains (JFH-1, a clinical isolate, and Jc1-n, a laboratory strain), which have different viral assembly and release characteristics. We found that 1.27% and 3.28% of JFH-1 and Jc1-n intracellular viral RNAs, respectively, are used for producing and releasing progeny virions. Analysis of the Malthusian parameter of the HCV genome (i.e., initial growth rate) and the number of *de novo* infections (i.e., initial transmissibility) suggests that the leaving-home strategy provides a higher level of initial transmission for Jc1-n, while, in contrast, the stay-at-home strategy provides a higher initial growth rate for JFH-1. Thus, theoretical-experimental analysis of viral dynamics enables us to better understand the proliferation strategies of viruses. Ours is the first study to analyze stay-leave trade-offs during the viral life cycle and their significance for viral proliferation.

## Introduction

Hepatitis C virus (HCV) is an RNA virus specifically infecting liver cells. HCV produces progeny viruses rapidly, with ∼10^12^ copies sometimes observed in patients [1]. Following virus entry into target cells, viral genomic RNA produces structural proteins (S) and non-structural proteins (NS) (**Fig. 1A**). Using the genomic RNA as a template, the viral non-structural proteins amplify HCV RNA (“RNA replication”). Genomic RNA can also be assembled with viral structural proteins into progeny virions to be egressed outside of cells, creating the opportunity for transmission (in this study, we call the process including particle assembly and egress “release”). Thus, a single HCV genomic RNA molecule can be used either for RNA replication or for release, and the balance between these processes governs viral proliferation. The molecular mechanisms underlying each event in the viral life cycle have been extensively investigated [2, 3], yet the replication-release trade-off and its significance for viral proliferation remain poorly understood.

**Figure 1.**
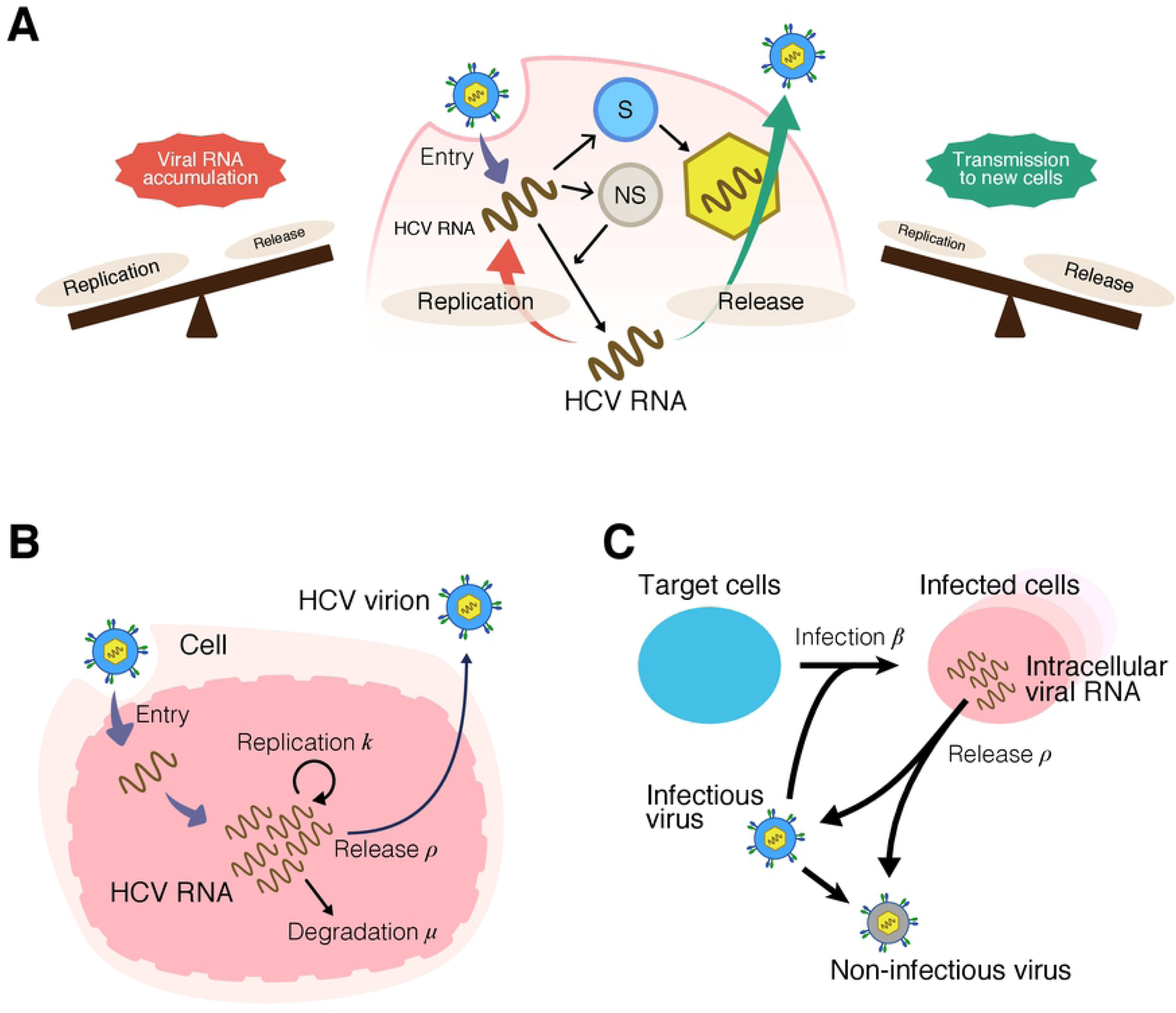
Schematic representation of multiscale HCV infection and mathematical model: **(A)** Schematic representation of intracellular HCV life cycle and trade-off between viral replication and release of intracellular viral RNA. Viral RNA in cells is translated to produce structural (S) and non-structural (NS) proteins. Viral RNA is either amplified through the functions of NS proteins through replication, or is assembled with S proteins and released as a progeny virus. If the balance between viral replication and release leans toward replication, intracellular viral RNAs will accumulate. In contrast, high rates of intracellular RNA release will create opportunities for transmission to new cells but will deplete viral RNA in the cell. **(B)** Modeling the intracellular virus life cycle. Intracellular viral RNA either replicates inside the cell at rate *k*, is degraded at rate *μ*, or assembles with viral proteins to be released within HCV virions at rate *ρ*. **(C)** Multiscale modeling of intracellular replication and intercellular infection. Target cells are infected by infectious viruses at rate *β*.

HCV JFH-1 is a genotype 2a strain isolated by our group from a patient with fulminant hepatitis [4]. JFH-1 has been a standard strain used for experiments to characterize HCV infection, virus-host interactions, and immune responses against HCV [4]. In addition, Jc1 or J6/JFH (a chimeric strain in which a region of the JFH-1 genome from the core to NS2 was replaced by sequences from another genotype 2a virus, the J6 strain) was developed as a laboratory strain to improve virus production, and used for development of antiviral agents and vaccines, which requires large amounts of virus [5, 6]. In spite of their high sequence similarity (97% identity over the whole genome), these two viruses have different virological characteristics especially in terms of the release process: while JFH-1 particles assemble on lipid droplet membranes, particle assembly of J6/JFH-1-chimeric lab strains is associated with endoplasmic reticulum-derived membranes [2, 3]. Thus, these two related strains are a useful a reference set to compare the dynamics of release and RNA replication.

In this study, we used a cell culture model of infection with these two HCV reference strains and measured the time-course of viral production (including HCV RNA inside cells and virions produced outside of the cells), infectivity of progeny HCV, and infected cell numbers. We also developed a multiscale mathematical model to describe intra- and inter-cellular HCV dynamics. This interdisciplinary approach suggests that different strategies exist for viral proliferation: the stay-at-home strategy (JFH-1) and the leaving-home strategy (Jc1-n, a J6/JFH-1-chimeric strain). We discuss the relevance of these strategies for viral proliferation, while referring to [7] for wider evolutionary context.

## Results

### Age-structured multiscale modeling of HCV infection

To describe the intracellular replication dynamics of HCV viral RNA, we used the following mathematical model:

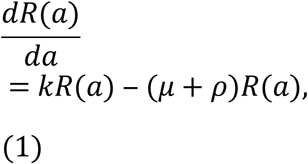

Where *R*(*a*) is the amount of intracellular viral RNA in cells that have been infected for time *a*. The intracellular viral RNA replicates at rate *k*, degrades at rate *μ*, and is released to extracellular space at rate *ρ* (**Fig. 1B**). Note that if viruses have small or large *ρ*, then they tend to stay inside or leave the cell, respectively (see later). In our virus experiments (see **Methods**), the released viruses could infect other target cells. To describe multi-round virus transmission (i.e., *de novo* infection), we needed to couple intracellular viral replication with a standard mathematical model for intercellular virus infection in cell culture [8, 9] (**Fig. 1C**). In **Supplementary Note 1**, we derived the following multiscale ordinary differential equation (ODE) model for HCV infection from the corresponding age-structured partial differential equation (PDE) model [10, 11]:

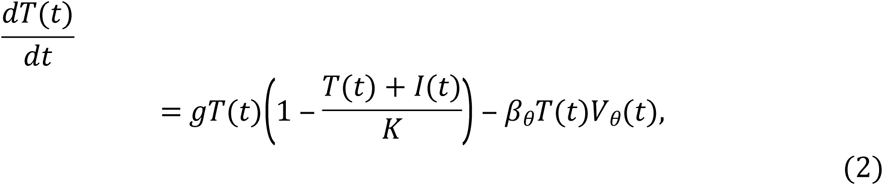

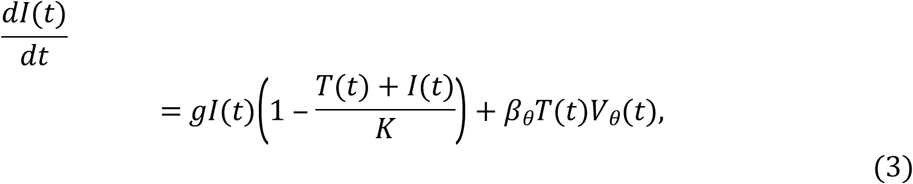

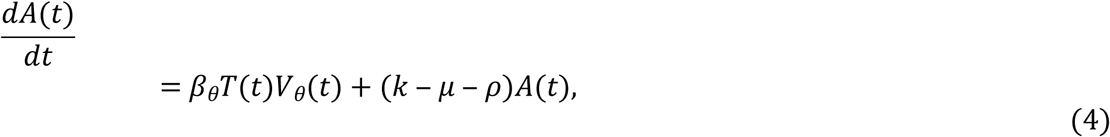

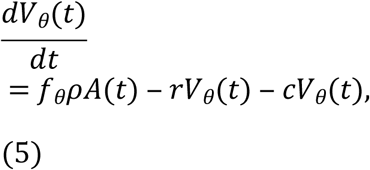

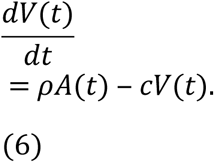

Here, the intercellular variables *T*(*t*) and *I*(*t*) represent the numbers of uninfected and infected target cells, and *V*(*t*) and *V*_*θ*_(*t*) represent the total amount of extracellular viral RNA (copies/well) and the extracellular viral infectious titer expressed (ffu/well), respectively. The intracellular variable *A*(*t*) represents the total amount of intracellular viral RNA. The parameters *g* and *K* represent the growth rate and the carrying capacity of target cells, and *β*_*θ*_ and *f*_*θ*_ are the converted infection rate constant and the fraction of infectious virus, respectively. We assumed that progeny viruses were cleared at rate *c*, and that infectious virions lose infectivity at rate *r*. Separately, we directly estimated *g, K, c, μ* + *ρ* and *r* for both HCV JFH-1 and Jc1-n in **Fig. S1**. Detailed explanations of Eqs. (2–6) are given in **Supplementary Note 1**.

To assess the variability of kinetic parameters and model predictions, we performed Bayesian estimation for the whole dataset using Markov chain Monte Carlo (MCMC) sampling (see **Methods**). Simultaneously, we fitted Eqs. (2–6) to the experimentally-determined numbers of uninfected and infected cells, extracellular viral RNA (copies/well) and infectious titer (ffu/well), and intracellular viral RNA (copies/well). These figures were derived from infection experiments using different numbers of plated cells for either HCV JFH-1 or Jc1-n as described previously [8, 9, 12, 13]. The remaining free model parameters (i.e., *β*_*θ*_, *k, ρ, f*_*θ*_) along with initial values for variables (i.e., *T*(0), *I*(0), *A*(0), *V*_*θ*_(*t*), *V*(0)) were determined. Experimental measurements below the detection limit were excluded in the fitting. The estimated parameters and initial values are listed in **Table 1** and **Table S1**. The typical behavior of the model using these best-fit parameter estimates is shown together with the data in **Fig. 2A** for HCV JFH-1 (orange) and Jc1-n (green) (see **Methods** for HCV strains), and indicated that Eqs. (2–6) described the *in vitro* data very well. The shadowed regions corresponded to 95% posterior predictive intervals, the solid and dashed lines gave the best-fit solution (mean) for Eqs. (2–6), and the orange circles and green triangles showed the experimental datasets.

**Table 1.** Parameter values estimated from the cell culture infection experiment.

| Parameter Name | Symbol | Unit | HCV JFH-1 |  | HCV Jc1 |  |
| --- | --- | --- | --- | --- | --- | --- |
|  |  |  | Value | 95% CI | Value | 95% CI |
| Fitted parameters from separate experiments |  |  |  |  |  |  |
| Rate of virion infectivity loss | $r$ | day<sup>-1</sup> | 1.60 | — | 2.40 | — |
| Degradation rate of extracellular viral RNA | $c_{RNA}$ | day<sup>-1</sup> | 0.08 | — | 0.24 | — |
| Clearance rate of extracellular viral RNA | $c_w$ | day<sup>-1</sup> | 1.18 | — | 1.82 | — |
| Reduction rate of intracellular viral RNA | $\mu + \rho$ | day<sup>-1</sup> | 0.80 | — | 0.89 | — |
| Estimated parameters from *in vitro* total cell growth data |  |  |  |  |  |  |
| Proliferation rate of Huh-7 cells | $g$ | day<sup>-1</sup> | 0.67 | — | 0.67 | — |
| Carrying capacity of Huh-7 cells | $K$ | cells | $4.12 \times 10^4$ | — | $3.75 \times 10^4$ | — |
| Parameters obtained from simultaneous fitting of full *in vitro* dataset |  |  |  |  |  |  |
| Rate constant for infections | $\beta_\theta$ | (ffu/well · day)<sup>-1</sup> | $1.29 \times 10^{-4}$ | $0.81\text{--}1.92 \times 10^{-4}$ | $2.21 \times 10^{-4}$ | $1.69\text{--}2.77 \times 10^{-4}$ |
| Replication rate of intracellular viral RNA | $k$ | day<sup>-1</sup> | 1.91 | 1.84–1.98 | 1.91 | 1.84–1.98 |
| Release rate of intracellular viral RNA | $\rho$ | day<sup>-1</sup> | $2.43 \times 10^{-2}$ | $1.87\text{--}3.11 \times 10^{-2}$ | $6.25 \times 10^{-2}$ | $4.62\text{--}8.44 \times 10^{-2}$ |
| Degradation rate of intracellular viral RNA | $\mu$ | day<sup>-1</sup> | 0.78 | 0.77–0.78 | 0.83 | 0.80–0.84 |
| Converted fraction of infectious viral RNA | $f_\theta$ | RNA copies · ffu<sup>-1</sup> | $1.21 \times 10^{-3}$ | $0.86\text{--}1.67 \times 10^{-3}$ | $1.13 \times 10^{-3}$ | $0.83\text{--}1.49 \times 10^{-3}$ |

**Figure 2.**
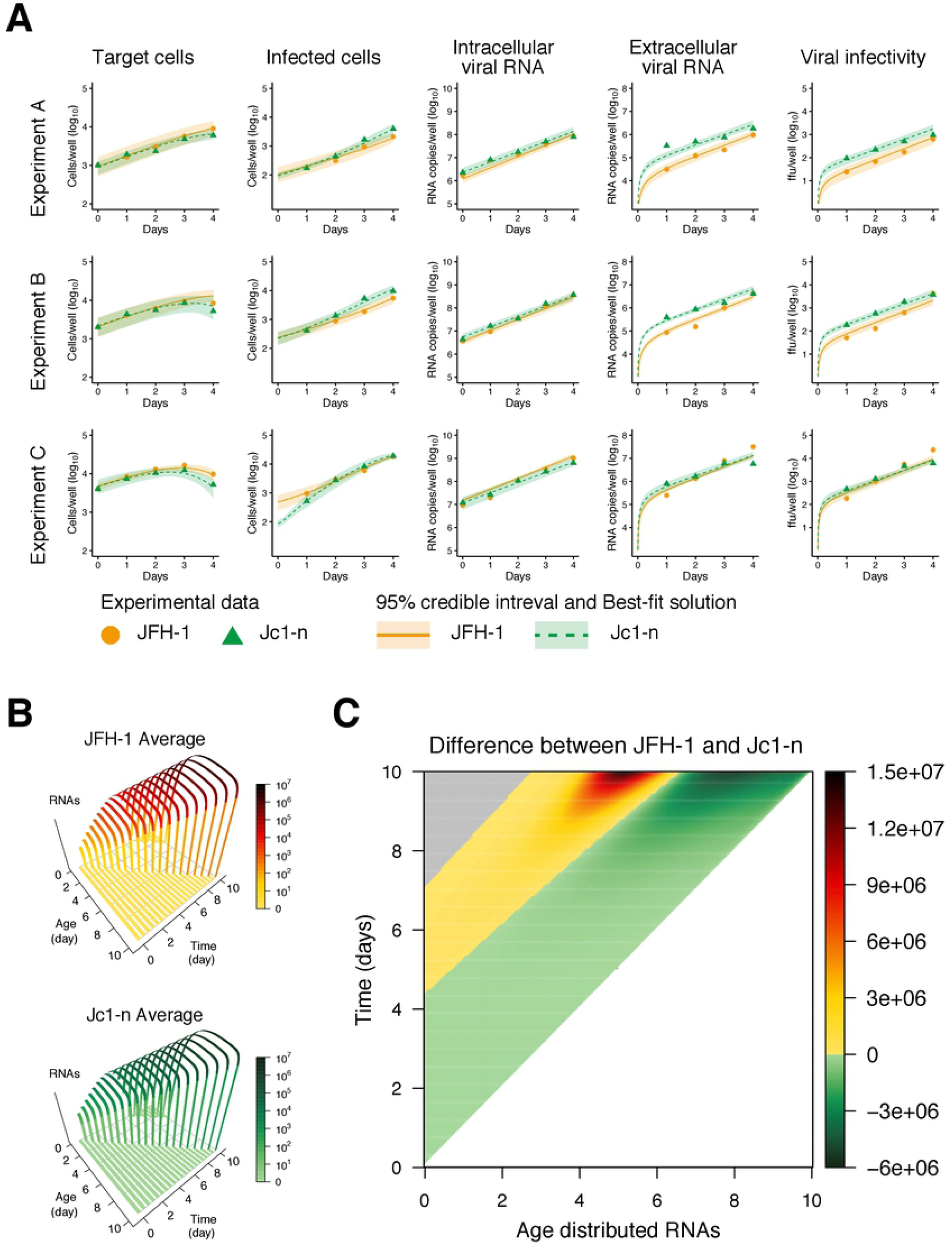
Dynamics of HCV JFH-1 and Jc1-n infection in cell culture. **(A)** Fitting of the mathematical model to the experimental data of HCV JFH-1 and Jc1-n infection in cell culture. Three different numbers of Huh-7 cells infected with either HCV JFH-1 or Jc1-n 1 day after inoculation were seeded (Experiment A: 1000, Experiment B: 2000, and Experiment C: 4000 cells per well) and chased to detect the following values at days 0, 1, 2, 3, and 4 post-seeding (log_10_ scale): numbers of uninfected and infected cells, amount of intracellular and extracellular viral RNA (copies/well), and extracellular viral infectivity (ffu/well) (orange circle: JFH-1, green triangle: Jc1-n). The shadowed regions correspond to 95% posterior intervals and the solid curves give the best-fit solution (mean) for Eqs. (2–6) to the time-course dataset. All data for each strain were fitted simultaneously. **(B)** Dynamics of the distributions of intracellular viral RNA according to infection age, *a*. The distributions were calculated using the original multiscale PDE model (Eqs. (S2–6) in **Supplementary Note 1**) using means of estimated parameters for HCV JFH-1 and Jc1-n. The colored bars represent the amount of intracellular viral RNA. **(C)** Difference in the distributions of intracellular viral RNA in total infected cells of infection age, *a*, between HCV JFH-1 and Jc1-n. The colored bar shows the difference in the amount of intracellular viral RNA (green: intracellular viral RNA during Jc1-n infection is more abundant than during JFH-1 infection, yellow-red-brown: intracellular viral RNA is more abundant for JFH-1 than for Jc1-n, gray: no new infection occurs due to depletion of target cells).

Using the estimated parameters shared between the original PDE model in **Supplementary Note 1** and the transformed ODE model (i.e., Eqs. (2–6)), we successfully reconstructed age information for intracellular viral RNA in infected cells of infection age *a*, which cannot be obtained through conventional experiments alone (**Fig. 2B**). **Fig. 2C** shows the differences in intracellular JFH-1 and Jc1-n viral RNA levels in cells of infection age *a*. At the beginning of the experiment, intracellular viral RNA increased faster under Jc1-n infection than under JFH-1 infection (shown in green). However, intracellular JFH-1 viral RNA gradually accumulated to higher levels than Jc1-n at later time points after infection (shown in yellow to brown). These data illustrated the different dynamics of these two strains and the impact of these dynamics on intracellular viral RNA production, all resulting from different strategies to transmit the viral genome (see below).

### Dynamics of HCV JFH-1 and Jc1-n strain replication

Our model [Eqs. (2–6)] applied to time-course experimental data allowed us to extract the following kinetic parameters: the distribution of the rate constant for infection, *β*_*θ*_, the release rate of intracellular viral RNA, *ρ*, the degradation rate of intracellular viral RNA, *μ*, the converted fraction of infectious viral RNA, *f*_*θ*_, and the replication rate of intracellular viral RNA, *k* (**Fig. 3** and **Table 1**). Comparing these parameters for JFH-1 and Jc1-n showed a significant difference between the rate constant for infections, *β*_*θ*_, of JFH-1 (1.29 × 10 ^‒ 4^, 95% CI: 0.81 ‒ 1.92 × 10 ^‒4^) and Jc1-n (2.21 × 10 ^‒ 4^, 95% CI: 1.69 ‒ 2.77 × 10 ^‒ 4^) (*p =* 1.82 × 10 ^‒ 4^ by repeated bootstrap *t*-test) (**Fig. 3A**). In addition, the release rate of intracellular viral RNA, *ρ*, for JFH-1 and Jc1-n were 2.43 × 10 ^‒ 2^ (95% CI: 1.87 ‒ 3.11 × 10 ^‒ 2^) and 6.25 × 10‒2 (95% CI: 4.62 ‒ 8.44 × 10 ^‒ 2^), respectively (*p =* 2.00 × 10 ^‒ 6^ by repeated bootstrap *t*-test) (**Fig. 3B**). These estimates indicated that Jc1-n infects cells 1.71 times faster and produces progeny viruses from infected cells 2.57 times faster than JFH-1. The estimate was further validated by independent experiments, in which Jc1-n entry and virus production were indeed significantly higher than those of JFH-1 (**Supplementary Note 2** and **Fig. S2**). There was also a small but significant difference between the degradation rate, *μ*, of JFH-1 (0.78, 95% CI: 0.77 ‒ 0.78) and Jc1-n (0.83, 95% CI: 0.80 ‒ 0.84) (**Fig. 3C**). No significant difference was apparent in the converted fraction of infectious virus, *f*_*θ*_ (**Fig. 3D**). Because JFH-1 and Jc1-n have identical non-structural regions essential for RNA replication (NS3–NS5B), we estimated the same viral RNA replication rate, *k*, for these two viruses (**Fig. 3E**). Hence, our parameter estimation captured the characteristics of the two strains well and was able to quantitatively describe viral infection dynamics.

**Figure 3.**
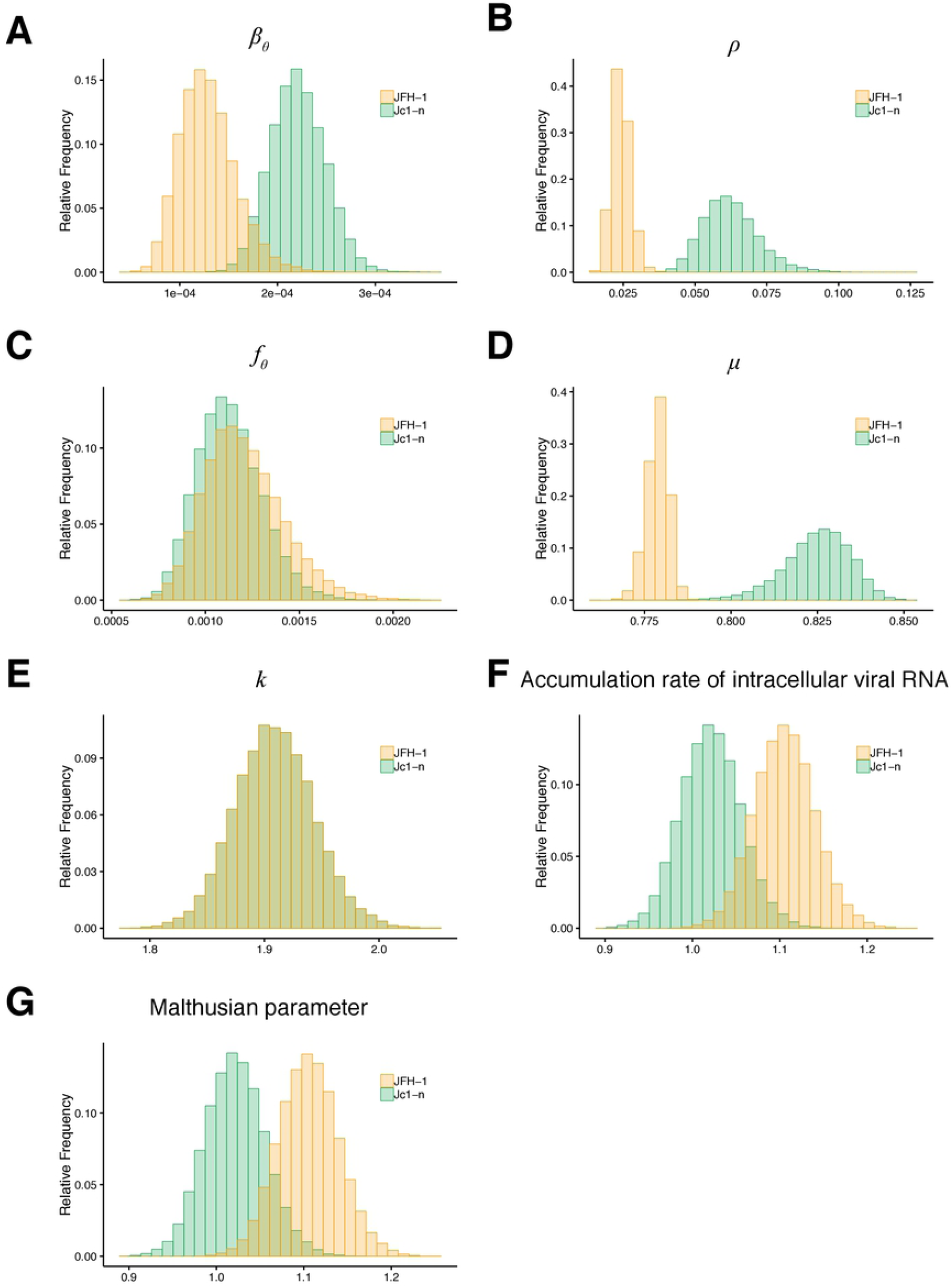
Characterization of viral dynamics of HCV JFH-1 and Jc1-n. The distributions of the rate constant for infection, *β*_*θ*_, the release rate of intracellular viral RNA, *ρ*, the degradation rate of intracellular viral RNA, *μ*, the converted fraction of infectious viral RNA, *f*_*θ*_, and the replication rate of intracellular viral RNA, *k*, inferred by MCMC computations are shown in **(A), (B), (C), (D)** and **(E)**, respectively, for HCV JFH-1 (orange) and Jc1-n (green). Parameters *β*_*θ*_, *ρ* and *μ* for Jc1-n were significantly larger than for JFH-1, while there was no significant difference in *f*_*θ*_ between the two strains as assessed by repeated bootstrap *t*-test. JFH-1 and Jc1-n stains had identical viral RNA replication rates. The distributions of accumulation rates of intracellular viral RNA, *k* ‒ *μ* ‒ *ρ*, and the Malthusian parameter, *M*, calculated from all accepted MCMC parameter estimates are shown in **(F)** and **(G)**, respectively, for HCV JFH-1 (orange) and Jc1-n (green). These indices were significantly larger for JFH-1 than for Jc1-n as assessed by the repeated bootstrap *t*-test.

In our multiscale model (Eqs. (2–6)), the accumulation rate of intracellular viral RNA was defined as the difference between the replication rate and the sum of the degradation rate and the release rate (i.e., *k* ‒ *μ* ‒ *ρ*). The distributions of calculated intracellular RNA accumulation rates for JFH-1 (1.11, 95% CI: 1.04 ‒ 1.18) and Jc1-n (1.02, 95% CI: 0.95 ‒ 1.09) are shown in **Fig. 3F** (*p =* 1.58 × 10 ^‒ 3^ by bootstrap *t*-test) (**Table 1**). The preferential accumulation of JFH-1 RNA inside cells was consistent with its tendency toward gradual increased levels of intracellular RNA at later time points (**Fig. 2C)**. To further evaluate total viral RNA level considering multi-round virus transmission, the Malthusian parameter, *M*, was used as an indicator of the initial growth rate of intracellular viral RNA for each HCV strain [8, 12, 14]. Here, the Malthusian parameter was given by

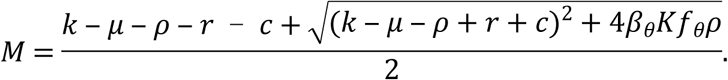

The Malthusian parameters for JFH-1 and Jc1-n were calculated as 1.11 (95% CI: 1.04 ‒ 1.18) and 1.02 (95% CI: 0.95 ‒ 1.09), respectively, and were significantly different from one another (*p =* 1.02 × 10 ^‒ 3^ by bootstrap *t*-test) (**Fig. 3G** and **Table 1**). Interestingly, even if Jc1-n had a larger infection rate, *β*_*θ*_, and release rate, *ρ*, compared with JFH-1, the initial growth rate of total JFH-1 RNA was higher than that of Jc1-n. This result demonstrated that the capacity to accumulate viral RNA inside cells predominantly determines the initial growth rate rather than release of progeny viruses.

### Stay-at-home strategy or leaving-home strategy for “optimizing” HCV proliferation

We investigated how differences between the two strains, JFH-1 and Jc1-n, might be interpreted in an evolutionary perspective. As mentioned above, we considered two opposing strategies: the “stay-at-home strategy” and the “leaving-home strategy”: If viruses have smaller *ρ*, they preferentially stay inside the cell, but if they have larger *ρ*, they leave the cell. To quantitatively characterize these different strategies, we defined the fraction of viral RNA remaining in the cells ((*k* ‒ *μ* ‒ *ρ*)*/k*), released from the cells (*ρ/k*), and degraded in the cells (*μ/k*) within the total intracellular viral RNA produced (**Fig. 4A**). Using all accepted MCMC parameter estimates from the time-course experimental datasets, we calculated that the fractions of viral RNA remaining were 57.9% and 53.5%, the fractions of viral RNA degraded were 40.8% and 43.2%, and the fractions of viral RNA released were 1.27% and 3.28% for JFH-1 and Jc1-n, respectively (**Fig. 4B**). Notably, Jc1-n used intracellular viral RNA for virus release 2.58 times faster than JFH-1, explaining the rapid transmission of Jc1-n (**Fig. 2C**). These results indicate the preferential “leaving-home” strategy of Jc1-n as compared with JFH-1, which adopts a “stay-at-home” strategy.

**Figure 4.**
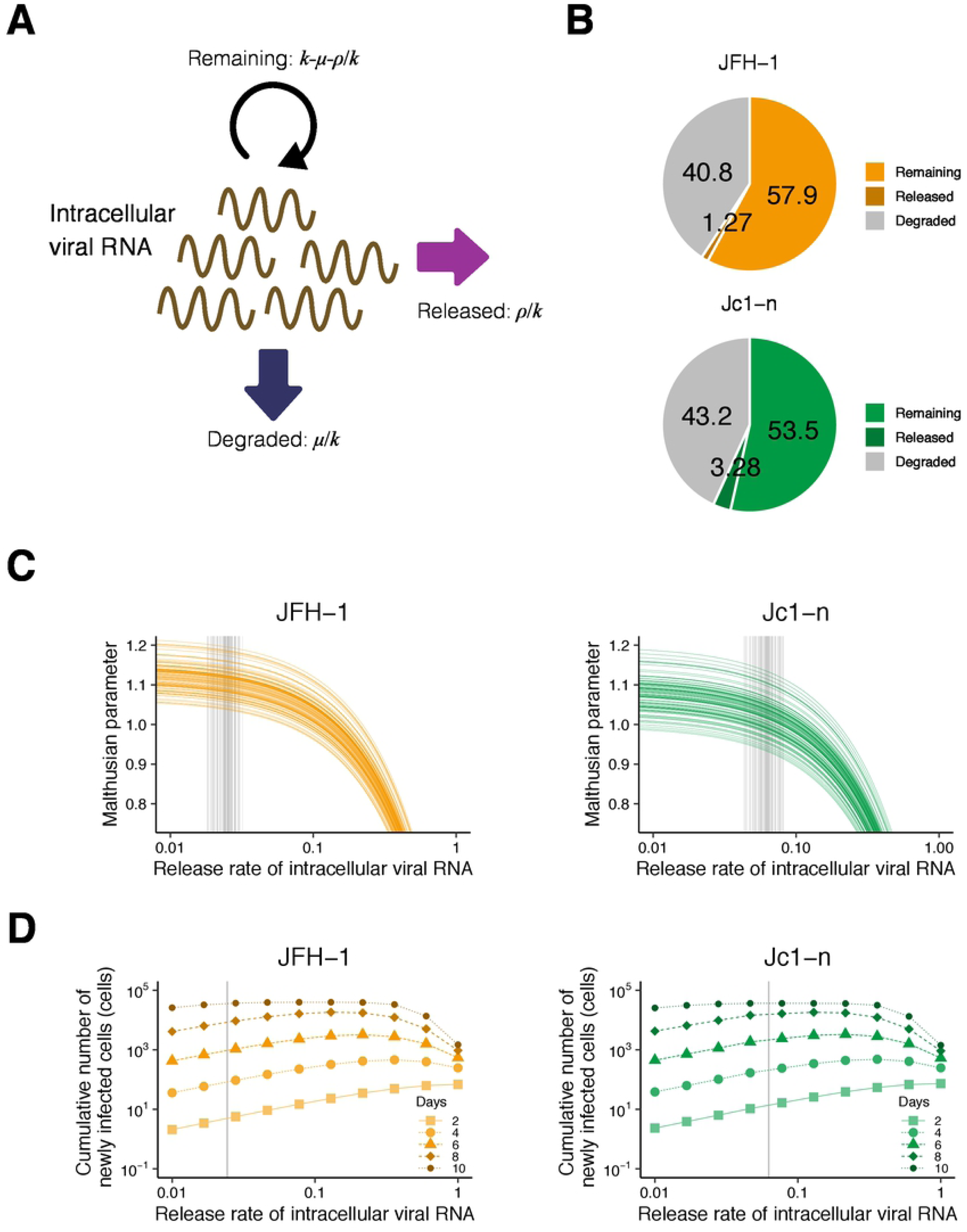
Different strategies adopted by JFH-1 and Jc1-n for viral proliferation. **(A)** Schematic representation of the fate of replicated intracellular viral RNA. Viral RNA is used either for driving RNA replication in cells, for producing progeny viruses for release outside cells, or is degraded. **(B)** Percentage of replicated intracellular HCV JFH-1 and Jc1-n viral RNA that remains inside cells, is released outside cells, and is degraded. **(C)** Change in the Malthusian parameter with various release rates of intracellular viral RNA. The orange and green curves show Malthusian parameters calculated using 100 parameter sets sampled from MCMC parameter estimates as functions of *ρ* for JFH-1 and Jc1-n, respectively. The gray vertical lines are the corresponding release rates estimated from the actual experimental data. **(D)** Change in the cumulative number of newly infected cells with the various release rates. The orange and green curves represent the cumulative number of newly infected cells until 2, 4, 6, 8, and 10 days post-infection calculated using the means of estimated parameters as function of *ρ* for JFH-1 and Jc1-n, respectively. The gray vertical line represents the mean release rate estimated from the experimental data.

To further investigate these two opposing strategies, we addressed the relevance of viral RNA release rates for viral proliferation using *in silico* analysis. With various values of the release rate of intracellular viral RNA, *ρ*, we calculated the Malthusian parameter for each strain as an indicator of viral fitness (**Fig. 4C**). Each curve shows Malthusian parameters calculated using 100 parameter sets sampled from MCMC parameter estimates as functions of *ρ*, and each gray vertical line is the corresponding estimated release rate. Interestingly, the smaller release rate, the larger the Malthusian parameter HCV achieves. This is because intracellular viral RNAs can be amplified faster compared with viral RNAs outside of cells that are degraded or enter new cells. This result showed that the JFH-1 strain is more optimized in terms of its Malthusian parameter compared with Jc1-n because of the smaller estimated values of *ρ*. That is, HCV JFH-1 adopts the stay-at-home strategy for acquiring a higher initial growth rate.

Next, we defined the cumulative number of newly infected cells at time evaluate viral transmissibility:

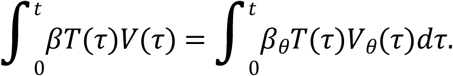

We also calculated the cumulative number of newly infected cells for each strain using the means of the estimated parameters as functions of *ρ* (**Fig. 4D**). Each curve showed the calculated cumulative number of infected cells until 2, 4, 6, 8, and 10 days post-infection, and the gray vertical line represented the mean release rate estimated from the infection experiment. The value of the release rate, which maximized the cumulative number of newly infected cells, was between 0.1 and 1. This is because an intermediate release rate effectively increases extracellular viral RNA for new infection: Lower release rates do not effectively produce new infections while higher release rates decrease intracellular viral RNA levels and thus diminish future new infections. Thus, it appears that Jc1-n is more optimized for producing newly infected cells. This implies that HCV Jc1-n adopts the leaving-home strategy to acquire an advantage in producing newly-infected cells.

Taken together, our theoretical investigation based on viral infection experiments revealed that the JFH-1 strain optimizes its initial growth rate, but the Jc1-n strain optimizes *de novo* infection. Ours is the first report to quantitatively evaluate these opposing evolutionary strategies and to show their significance for virus proliferation at the intracellular and intercellular levels.

## Discussion

Through a combined experimental-theoretical approach, we analyzed the dynamics of the HCV life cycle using two related HCV strains, JFH-1 and Jc1-n, employing different particle assembly/release strategies. We quantified the intra- and inter-cellular viral dynamics of these strains by applying an age-structured multiscale model to time-course experimental data from an HCV infection cell culture assay (**Fig. 2A** and **Table 1**): As in [10, 11], we transformed the multiscale model formulated by PDEs to an identical multiscale ODE model (i.e., Eqs. (2–6)), and estimated parameters shared between the PDE and ODE models. It is technically challenging to obtain experimental measurements with age information, but thanks to the estimated values of these common parameters, we managed to reconstruct age information for intracellular viral RNA (**Fig. 2BC**). In addition, comparing the calculated Malthusian parameters and the cumulative number of newly infected cells between the two strains (**Fig. 3FG**), we found that the JFH-1 strain had a higher initial growth rate but that Jc1 produced more *de novo* infections.

Based on our results, we propose two opposing strategies for viral proliferation: the “stay-at-home strategy” and the “leaving-home strategy.” From an evolutionary perspective, JFH-1 adopts a stay-at-home strategy and preferentially uses viral genomic RNA for increasing intracellular replication. In contrast, adopting a leaving-home strategy, Jc1-n uses more viral genomic RNA for producing progeny virions capable of new transmission events to increase the number of infected cells (**Fig. 4**). Thus, Jc1-n infects cells 1.71 times faster and produces viral RNA from infected cells 2.57 times faster than JFH-1. Our group and others reported that JFH-1 assembled progeny virions on the membranes of hepatic lipid droplets, while J6/JFH-1 chimeric strains mainly used endoplasmic reticulum-derived membranes for particle production [2, 3]. Although the molecular aspects of this difference have been analyzed, its significance for viral proliferation and dynamics is not completely understood. Our results raise the possibility that different subcellular locations for particle assembly impact the rates of particle assembly and release, which in turn determine virus proliferation. Further analysis might shed light on why one HCV strain has to be assembled on the lipid droplet membrane while another assembles in association with the endoplasmic reticulum.

The choice of replication strategy not only determines virus proliferation but also affects the pathogenic features of the virus: JFH-1, which preferentially amplifies intracellular RNA, caused fulminant hepatitis with rapid viral replication and severe inflammation. By contrast, J6, the original strain encoding the Jc1-n structural region, was isolated from a patient with chronic hepatitis and generally replicates more moderately, with robust spread of infected cells used as a longer term strategy to establish persistent infection. Characterization of the proliferation strategies of viruses is of significant importance when trying to understand their clinical as well as evolutionary properties.

## Methods

### Cell culture and HCV infection

Huh-7.5.1 (kindly provided by Dr. Francis Chisari, The Scripps Research Institute) and Huh7-25 cells were cultured in Dulbecco’s Modified Eagle’s Medium (Invitrogen) supplemented with 10% fetal bovine serum (Sigma), 10 units/mL penicillin, 10 mg/mL streptomycin, 0.1 mM non-essential amino acids (Invitrogen), 1 mM sodium pyruvate, and 10 mM HEPES, pH 7.4, at 37°C under a humidified atmosphere containing 5% CO_2_. We used HCV strains JFH-1, a genotype 2a clinical isolate from a patient with fulminant hepatitis [4], and Jc1-n, a J6/JFH-1 chimeric laboratory strain [13]. JFH-1 and Jc1-n have 96.7% amino acid identity over the whole genome. HCV inoculum for infection experiments was recovered from the culture supernatants of Huh-7.5.1 cells transfected with the corresponding HCV RNA as described [4]. Huh-7.5.1 cells were inoculated with JFH-1 or Jc1-n for 4 h and then passaged to seed a new 96 well plate at different cell densities (1000, 2000, or 4000 cells/well). At days 0, 1, 2, 3, and 4 post-seeding, culture supernatants and cell lysates were recovered to quantify HCV RNA by real time RT-PCR as previously described [13]. The infectivity of HCV in culture supernatants was measured using a focus-forming assay as described [13]. To quantify the number of uninfected and infected cells, cells were fixed and stained with anti-HCV core antibody by immunofluorescence assay as described [13].

### Data fitting and parameter estimation

The parameters *g* and *K* were separately estimated (see **Supplementary Note 3**) and fixed at 0.660 and 4.12 × 10^4^, respectively, for the JFH-1 strain, and 0.665 and 3.75 × 10^4^, respectively, for the Jc1-n strain. A statistical model adopted from Bayesian inference assumed that measurement error followed a normal distribution with mean zero and unknown variance (error variance). A distribution of error variance was also inferred using the Gamma distribution as its prior distribution. The posterior predictive parameter distribution as an output of MCMC computation represented parameter variability. Distributions of model parameters (i.e., *β*_*θ*_, *k, ρ, f*_*θ*_) and initial values (i.e., *T*(0), *I*(0), *A*(0), *V*_*θ*_(*t*), *V*(0)) in Eqs. (2–6) were inferred directly by MCMC computations. Distributions of derived quantities were calculated from the inferred parameter sets (**Fig. 3EF** for graphical representation). A set of computations for Eqs (2–6) with estimated parameter sets gives a distribution of outputs (the number of cells and the intra- and extra-cellular viral loads) as model predictions. To investigate variation in model predictions, global sensitivity analyses were performed. The range of possible variation is shown in **Fig. 2A** as 95% confidence intervals. Technical details of MCMC computations are summarized below.

### Statistical analysis

Package FME [15] in R Statistical Software [16] was used to infer posterior predictive parameter distributions. The delayed rejection and Metropolis method [17] was used as a default computation scheme for FME to perform MCMC computations. MCMC computations for parameter inference were implemented using the pre-defined function modMCMC() in package FME as introduced in **Methods**. Convergence of Markov chains to a stationary distribution was required to ensure parameter sets were sampled from a posterior distribution. Only the last 90000 of 100000 chains were used as burn-in. The convergence of the last 90000 chains was manually checked with figures produced by package coda [18], a collection of diagnostic tools for MCMC computation. The 95% credible interval shown as a shadowed region in each panel of **Fig. 2A** was produced from 100 randomly chosen inferred parameter sets and corresponding model predictions. We employed a bootstrap *t*-test [19] to quantitatively characterize differences in parameters and derived quantities between HCV JFH-1 and Jc1-n. In total, 100000 parameter sets were sampled with replacement from the posterior predictive distributions to calculate the bootstrap t-statistics. To avoid potential sampling bias, the bootstrap t-test was performed 100 times repeatedly. The averages of the computed p-values were used as indicators of differences.

## ACKNOWLEDGMENTS

Huh7.5.1 cells were kindly provided by Dr. Francis Chisari at The Scripps Research Institute. This study was supported in part by Grants-in-Aid for JSPS Research Fellow 19J12319 (to S. Iwanami), 19J21395 (to K.K.), Scientific Research (KAKENHI) Scientific Research B 18KT0018 (to S.I.), 18H01139 (to S.I.), 16H04845 (to S.I.), 17H04085 (to K.W.), Scientific Research in Innovative Areas 19H04839 (to S.I.), 18H05103 (to S.I.); AMED CREST 19gm1310002 (to S.I.); AMED J-PRIDE 19fm0208006s0103 (to S.I.), 19fm0208014h0003 (to S.I.), 19fm0208019h0103 (to S.I.), 19fm0208019j0003 (to K.W.); AMED Research Program on HIV/AIDS 19fk0410023s0101 (to S.I.); Research Program on Emerging and Re-emerging Infectious Diseases 19fk0108050h0003 (to S.I.); Program for Basic and Clinical Research on Hepatitis 19fk0210036h0502 (to S.I.), 19fk0210036j0002 (to K.W.); Program on the Innovative Development and the Application of New Drugs for Hepatitis B 19fk0310114h0103 (to S.I.), 19fk0310114j0003 (to K.W.), 19fk0310101j1003 (to K.W.), 19fk0310103j0203 (to K.W.); JST MIRAI (to S.I. and K.W.); JST CREST (to S.I. and K.W.); Mitsui Life Social Welfare Foundation (to S.I. and K.W.); Shin-Nihon of Advanced Medical Research (to S.I.); Suzuken Memorial Foundation (to S.I.); Life Science Foundation of Japan (to S.I.); SECOM Science and Technology Foundation (to S.I.); The Japan Prize Foundation (to S.I.); Toyota Physical and Chemical Research Institute (to S.I.); The Yasuda Medical Foundation (to K.W.); Smoking Research Foundation (to K.W.); Takeda Science Foundation (to K.W.).

## AUTHOR CONTRIBUTIONS

OD, S Iwami and KW designed the research. HO, KN, and KW conducted the experiments. S Iwanami, KK, YA, HI and S Iwami carried out the computational analysis. OD, S Iwami and KW supervised the project. All authors contributed to writing the manuscript.

